# The Zinc Finger Antiviral Protein restricts SARS-CoV-2

**DOI:** 10.1101/2020.06.04.134379

**Authors:** Rayhane Nchioua, Dorota Kmiec, Janis Müller, Carina Conzelmann, Rüdiger Groß, Chad Swanson, Stuart Neil, Steffen Stenger, Daniel Sauter, Jan Münch, Konstantin M. J. Sparrer, Frank Kirchhoff

## Abstract

Recent evidence shows that the Severe Acute Respiratory Syndrome Coronavirus 2 (SARS-CoV-2) is highly sensitive to interferons (IFNs). However, the underlying antiviral effectors remain to be defined. Here, we show that Zinc finger antiviral protein (ZAP) that specifically targets CpG dinucleotides in viral RNA sequences restricts SARS-CoV-2. We demonstrate that ZAP and its cofactors KHNYN and TRIM25 are expressed in human lung cells. Type I, II and III IFNs all strongly inhibited SARS-CoV-2 and further induced ZAP expression. Strikingly, SARS-CoV-2 and its closest relatives from bats show the strongest CpG suppression among all known human and bat coronaviruses, respectively. Nevertheless, knock-down of ZAP significantly increased SARS-CoV-2 production in lung cells, particularly upon treatment with IFN-α or IFN-γ. Thus, our results identify ZAP as an effector of the IFN response against SARS-CoV-2, although this pandemic pathogen may be preadapted to the low CpG environment in humans.

**Highlights:**

- SARS-CoV-2 and its closest bat relatives show strong CpG suppression
- IFN-β, -γ and -λ inhibit SARS-CoV-2 with high efficiency
- ZAP restricts SARS-CoV-2 and contributes to the antiviral effect of IFNs

## INTRODUCTION

SARS-CoV-2, the causative agent of coronavirus disease 2019 (COVID-19), has been first detected in humans in Wuhan China at the end of 2019 and rapidly spreads in human populations causing a devastating pandemic (Zhou et al., 2020). As of June 2020, almost 7 million infections with SARS-CoV-2 around the globe have been confirmed and the virus has caused about 395.000 deaths (https://coronavirus.jhu.edu/map.html). While SARS-CoV-2 usually causes no or relatively mild respiratory infections in younger individuals, it regularly results in severe respiratory disease and death in the elderly and in people with specific medical conditions, such as asthma, heart diseases, diabetes or severe obesity (Zheng et al., 2020). SARS-CoV-2 is spreading substantially more efficiently than the first zoonotic highly pathogenic coronavirus (SARS-CoV) that emerged in 2002 and infected about 8.000 individuals (Graham and Baric, 2010; Petrosillo et al., 2020). Despite its rapid global spread, SARS-CoV-2 seems to be more susceptible to inhibition by type I IFNs representing a major component of the first line of innate antiviral immune defence than SARS-CoV (Mantlo et al., 2020). Consequently, type I IFNs are currently considered for treatment of COVID-19 (Sallard et al., 2020).

Treatment with IFNs induces the expression of hundreds of cellular IFN-stimulated genes (ISGs), and it is currently unknown which of these genes contribute to IFN-inducible restriction of SARS-CoV-2 replication. However, antiviral factors may exert strong selection pressure and result in specific viral properties that provide hints for efficient IFN-mediated immune responses. For example, it is long known that coronaviruses display marked suppression of CpG dinucleotides (Woo et al., 2007) and recent evidence suggests that this is also the case for SARS-CoV-2 (Xia, 2020). At least in part, this CpG suppression may be driven by the zinc finger antiviral protein (ZAP) that restricts numerous viral pathogens (Ghimire et al., 2018) and specifically targets CpG-rich RNA sequences that are underrepresented in the human transcriptome (Takata et al., 2017).

Coronaviruses (CoVs) are found in numerous animal species, such as bats, swine, cattle, horses, camels, cats, dogs, rodents, rabbits, ferrets, civets, pangolins, birds and snakes (Corman et al., 2018; Cui et al., 2019). They have successfully crossed the species-barrier to humans at least seven times and it is thought that all human CoVs (hCoVs) originate from ancestral bat viruses, although intermediate hosts frequently facilitated viral zoonoses (Banerjee et al., 2019). Four human coronaviruses are associated with seasonal common colds. Two of these (CoV-229E and OC43) have been identified more than 60 years ago and are relatively well adapted to humans. Two other coronaviruses associated with a range of respiratory symptoms have been identified in 2004 (CoV-NL63) and 2005 (CoV-HKU1), respectively (Van Der Hoek et al., 2004; Woo et al., 2006). While these strains usually cause mild respiratory diseases, three additional coronaviruses responsible for severe lung disease emerged from viral zoonoses in the last twenty years. In 2003, SARS-CoV was identified as causative agent of severe acute respiratory syndromes (SARS) with ~10% mortality (Ksiazek et al., 2003). The highly lethal MERS-CoV appeared in 2012 and was associated with case-fatality rates of almost 40% (Bermingham et al., 2012). The current SARS-CoV-2 shows a lower case-fatality rate (~2%) but is spreading at enormous speed. While the direct animal precursor remains to be identified, close relatives of SARS-CoV-2 have been detected in bats (Zhou et al., 2020a, 2020b) and pangolins (Lam et al., 2020; Xiao et al., 2020).

To define selection pressures on SARS-CoV-2 and other coronaviruses, we examined CpG frequencies and distribution in all seven human viruses and their closest animal counterparts. We found that CpG dinucleotides are generally suppressed and observed a trend towards lower CpG frequencies in hCoVs compared to their non-human relatives. In agreement with recent data (MacLean et al., 2020; Xia, 2020), SARS-CoV-2 showed stronger CpG suppression than SARS-CoV and MERS-CoV, albeit with substantial variation across its genome (Digard et al., 2020). Remarkably, the closest bat relatives of SARS-CoV-2 display the strongest CpG suppression of all coronaviruses available from this natural reservoir host. Furthermore, we found that the CpG targeting host factor ZAP is expressed in human lung cells and restricts SARS-CoV-2 especially in the presence of IFNs. Our data suggest that zoonotic transmission of a coronavirus with an unusually low frequency of CpG dinucleotides facilitated the pandemic spread of SARS-CoV-2 although it does not confer full resistance to ZAP-mediated restriction.

## RESULTS

### SARS-CoV-2 and its closest bat relatives show unusually strong CpG suppression

To determine the frequency and distribution of CpG dinucleotides and to identify possible differences in the levels of suppression, we analysed 67 genomes representing the seven human coronaviruses (hCoVs) and their closest animal relatives (Table S1; Figure 1A). Direct animal precursors or close relatives of the emerging human SARS-, MERS-, SARS-CoV-2, as well as seasonal hCoV-229E and hCoV-OC43 coronaviruses have been previously identified (Figure 1A). In contrast, the closest known animal relatives of hCoV-HKU1 and hCoV-NL63 found in rats and bats show only ~74% sequence identity to the respective human coronaviruses (Table S1), indicating long evolutionary divergence (Dominguez et al., 2012). Even though the immediate animal precursors are not always known, it is assumed that all seven hCoVs originate from bats, mice or domestic animals, where bats that harbour an enormous diversity of CoVs represent the reservoir host (Cui et al., 2019; Ye et al., 2020).

**Figure 1:**
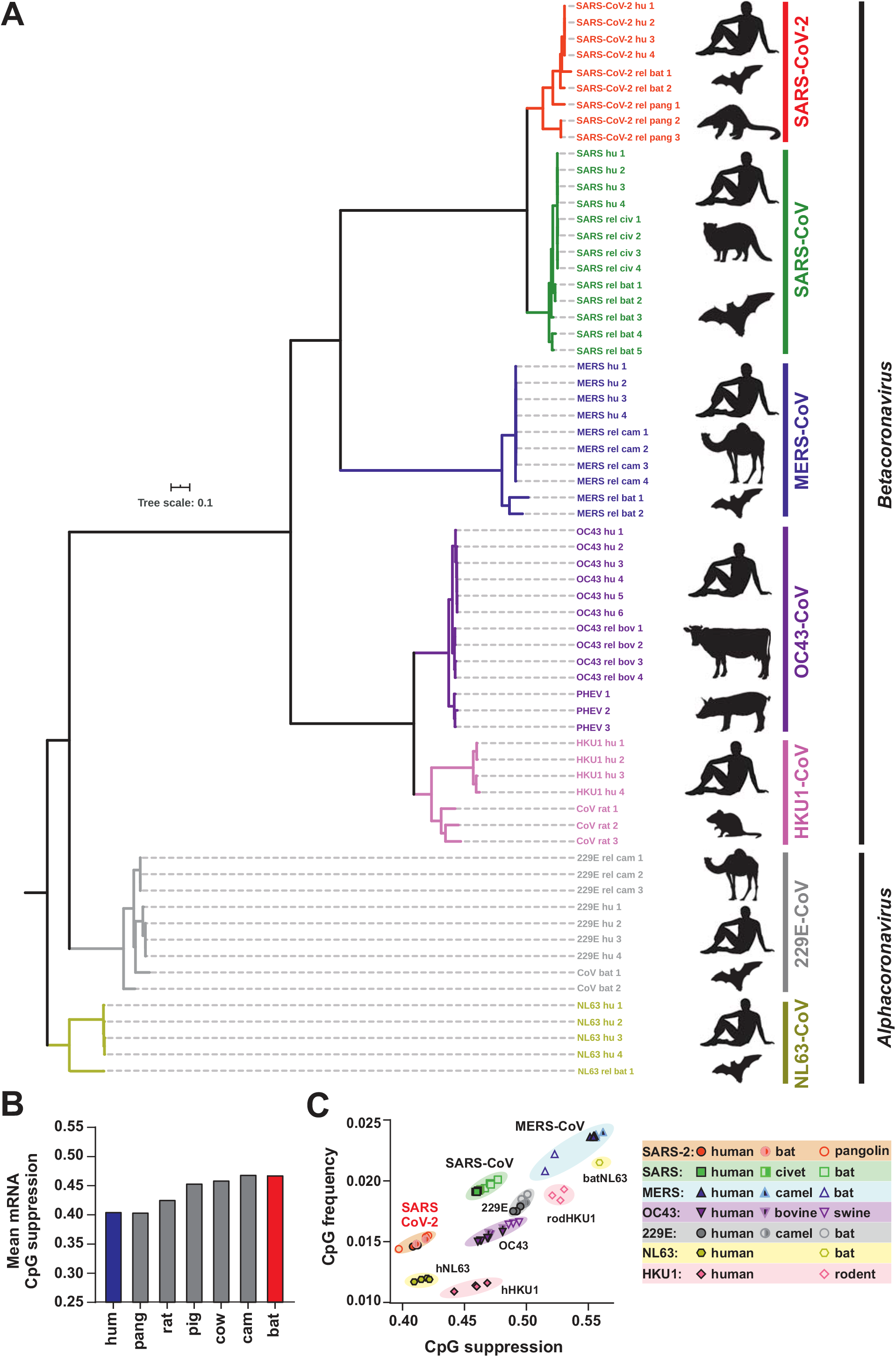
Phylogenetic relationship between human coronaviruses and their animal relatives and CpG suppression in pathogens and their hosts. **(A)** Distance-based relationship inference based on representative full genome nucleotide sequences of human SARS-CoV-2, SARS, MERS, HKU-1, OC43, NL63 and 229E strains and their closest known animal relatives. Black symbol to the right indicates the viral host (human, bat, pangolin, civet, camel, rat, pig, cattle). **(B)** Mean CpG suppression (i.e. number of observed CpGs normalized to expected CpGs based on sequence length and GC content) in mRNAs of the indicated host species (human, pangolin, rat, pig, cow, camel and bat). **(C)** CpG frequency (no. of CpGs normalized to sequence nucleotide length) and suppression (no. of observed CpGs normalized to expected CpGs based on sequence length and GC content) in human coronavirus genomes and their closest animal-infecting relatives. See also Table S1.

Vertebrate RNA viruses are known to mimic the CpG suppression of their hosts and increased viral CpG suppression following zoonotic transmission has been proposed to represent an important human-specific adaptation (Greenbaum et al., 2008). To assess whether zoonotic transmission of CoVs to humans might increase the selection pressure against CpGs, we first analysed the levels of CpG suppression in the reservoir bat, intermediate and human hosts. While the human genome and transcriptome has been extensively studied and undergone multiple quality checks, the transcript datasets of other species mostly contain predicted mRNA sequences and could be biased by the presence of poor-quality transcripts or modelling errors. We have therefore removed mRNA transcripts containing stretches of non-ATCG bases from the analysis and also quantified length-dependent CpG suppression to determine if differences in average suppression are consistent across datasets that included at least 40.000 transcripts of each species. Overall, the levels of CpG suppression vary and inversely correlate with the length of the cellular RNAs (Figure S1A), most likely due to the presence of regulatory elements in the 5’ UTR (Deaton and Bird, 2011; Saxonov et al., 2006). On average, however, CpG suppression is more pronounced in humans compared to bats, while the rat, pig and cow hosts show an intermediate phenotype and camels being similar to bats and pangolins to humans, respectively (Figure 1B). Analysis of the genomes of human coronaviruses and their animal relatives revealed that all of them show significant CpG suppression, although with varying extent (0.39-0.67; Figure 1C). MERS-CoV, associated with highest host mortality but also most limited spread, is the least CpG suppressed human coronavirus. In contrast, SARS-CoV-2 shows the strongest CpG suppression approximating the levels of suppression existing in its human host (Figure 1C; Table S1).

To assess whether the selection pressure against viral CpGs increases after zoonotic transmission, we compared CpG suppression and frequency, as well as GC content of hCoVs and their closest known animal relatives. Community-acquired hCoVs showed significantly lower CpG frequencies and stronger suppression than their closest animal relatives, while this was not the case for the highly pathogenic SARS- and MERS-CoVs (Figure 1C, 2A). SARS- and MERS-CoV show higher genomic GC content than the remaining CoVs, which explains why they display higher CpG frequencies at the same level of CpG suppression (Figure 2A). SARS-CoV-2 and its closest relatives from bats and pangolins show stronger CpG suppression than most other CoVs. Consequently, their CpG frequencies are similar to those found in community-acquired CoVs and lower than in SARS-CoV and MERS-CoV as well as their relatives, detected in bats, camels and civet cats (Figure 1C, 2A). These results raised the possibility that SARS-CoV-2 originated from a zoonotic virus showing a particularly low frequency of CpG dinucleotides. Indeed, the two closest animal relatives of SARS-CoV-2 (RaTG13 and RmYN02) show markedly lower CpG frequencies than all remaining 180 bat viruses available for analysis (Figure 2B).

**Figure 2:**
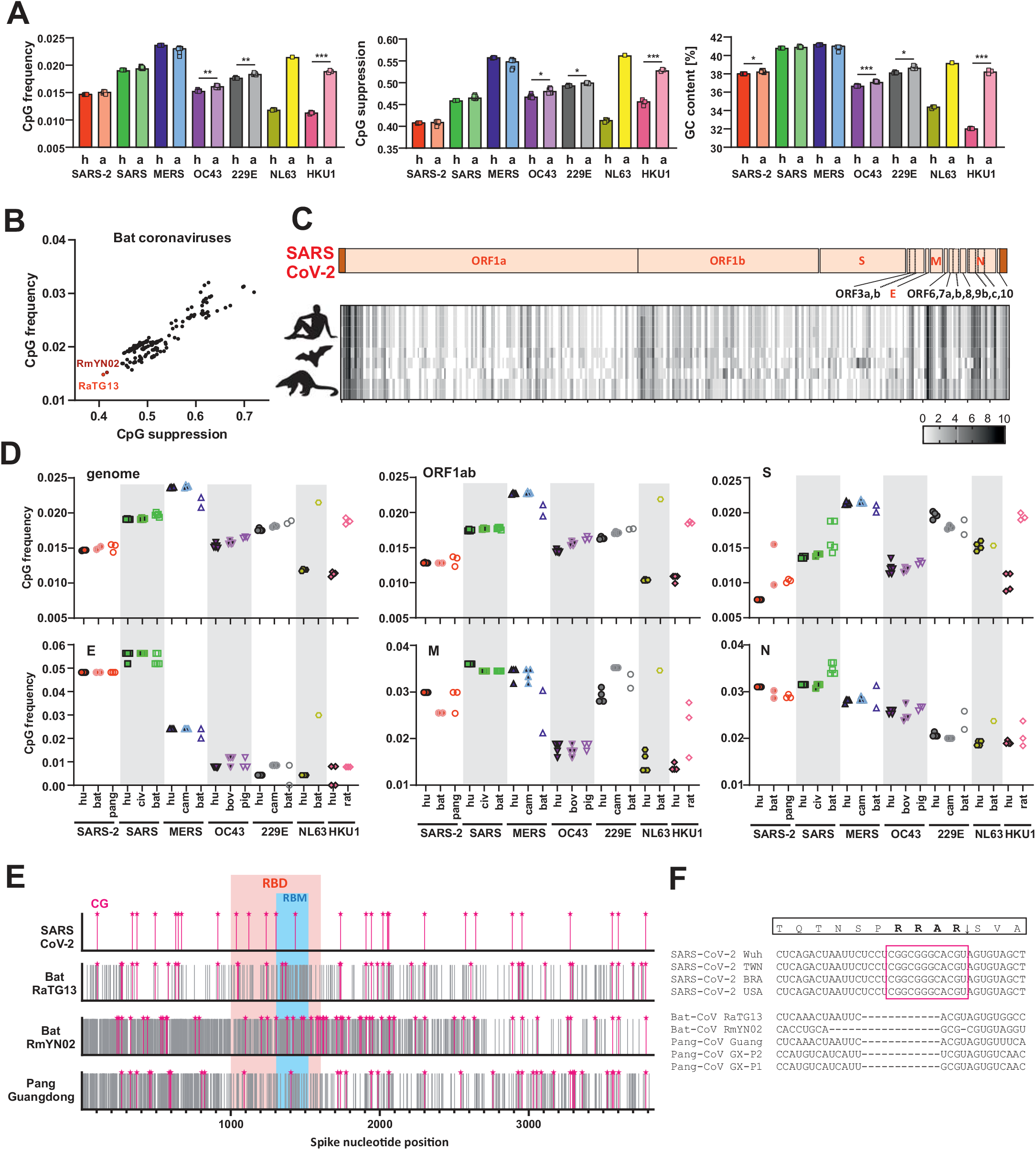
CpG content and distribution in human coronaviruses and their animal relatives. **(A)** CpG frequency, suppression and GC content in the genomes of human (h) and animal (a) infecting coronaviruses. **(B)** CpG frequency and suppression of sequenced bat coronaviruses available in NCBI database (n=182); close relatives of SARS-CoV-2 are shown in bright (RaTG13) and dark (RmYN02) red. Each point represents one viral strain. **(C)** Heatmap showing the number of CpG dinucleotides ranging from 0 (white) to 10 (black) within 100 bp sliding windows of aligned genomic sequences of SARS-CoV-2 and its closest relatives infecting bats and pangolins. Genome organisation diagram above represents that of the human virus. **(D)** CpG frequency in major genes of human and related animal coronaviruses. These genes encode viral polyproteins (ORF1ab), envelope (E), spike (S), nucleocapsid (N) and matrix (M) proteins. **(E)** Schematic representation of spike nucleotide sequences of SARS-CoV-2 and its closest relatives showing the relative position of CpGs (pink stars), receptor binding domain (RBD) and motif (RBM). Grey lines indicate nucleotide mismatches compared to aligned SARS-CoV-2 spike. **(F)** Insertion in spike of SARS-CoV-2 introducing a novel furin-cleavage site after an RRAR motif. See also Figure S1.

Coronavirus genomes differ in length and the presence of specific accessory genes (Table S1). Thus, we generated individual CpG distribution heatmaps for each group of hCoVs and their animal counterparts (Figures 2C, S1B) and compared CpG frequencies in the major viral genes (Figure 2D). On average, SARS-CoV-2 shows substantially lower CpG frequencies (0.014) than SARS-CoV (0.019) and MERS-CoV (0.024) (Figure 2A). However, we observed fluctuation between individual genes. While CpGs are strongly suppressed in the large ORF1a/b and Spike (S) ORFs, both SARS-CoV and SARS-CoV-2 show high numbers of CpGs in the 3’ regions of their genomes (Figure 2C, S1B). Consequently, they display higher CpG frequencies than other CoVs in the E (envelope) and (to a lesser extent) N (Nucleocapsid) coding regions (Figure 2D). However, the E gene encompasses just 228 to 267 bp. Thus, small changes in CpG numbers have a large impact on their frequency.

Notably, a region in the Spike gene of the bat CoV-RmYN02 strain that is otherwise closely related to SARS-CoV-2 (Zhou et al., 2020a) encoding amino acid residues involved in interaction with the viral ACE2 receptor shows low nucleotide identity and much higher frequency of CpGs than SARS-CoV-2 (Figure 2E). In addition, a small insertion that is present in SARS-CoV-2 Spike but not in its relatives from bats and pangolins not only introduced a potential furin cleavage site but also an additional clustered CpG motif that may be targeted by ZAP (Figure 2F).

Altogether, our results support that the selective pressure against CpGs is increased upon zoonotic transmission from bats and most intermediate hosts to humans. This indicates that the differences between hCoVs and their animal relatives may reflect different degrees of adaptation. At least in part, however, SARS-CoV-2 may already have been preadapted to the low CpG environment in humans because it’s closest known counterparts from bats contains an unusually low frequency of CpG dinucleotides.

### All three types of IFN inhibit SARS-CoV-2 and induce the short (S) isoform of ZAP

Our sequence analyses indicated that successful zoonotic transmission of CoVs to humans is associated with increased selection pressure for CpG suppression. To assess whether the antiviral factor ZAP might be the driving force behind this, we first examined whether ZAP is expressed in viral target cells. Western blot analyses of the human epithelial lung cancer cell lines Calu-3 and A549 that are commonly used in SARS-CoV-2 research (Hoffmann et al., 2020a; Matsuyama et al., 2020), as well as primary human lung fibroblasts, showed that all of these constitutively express the short and long isoforms of ZAP (Figure S2A-C). Treatment with TNFα as well as IFN-α, -β and -γ had modest effects on expression of the long isoform of ZAP but usually enhanced expression of the short isoform. IFN-γ had the most striking effects and increased ZAP(S) up to 8-fold (Figure S2, right panels). We also examined expression of TRIM25 and KHNYN because ZAP itself does not possess RNAse activity and it has been reported that these cofactors are critical for effective viral restriction (Li et al., 2017; Zheng et al., 2017; Ficarelli et al., 2019). TRIM25 and KHNYN were constitutively expressed in Calu-3 and A549 cells and the former is further induced by IFNs. Only marginal levels of KHNYN expression were detected in primary lung fibroblasts (Figure S2C).

IFNs are currently evaluated for the treatment of COVID-19 (Sallard et al., 2020). However, it is under debate which type of IFN is most effective against SARS-CoV-2 (Park and Iwasaki, 2020). To determine which IFNs are most potent in inhibiting SARS-CoV-2 and in inducing ZAP, we performed titration experiments using type I (α, β), II (γ) and III (λ) IFNs. We selected Calu-3 cells for these experiments because they are highly susceptible to SARS-CoV-2 infection (Chu et al., 2020; Hoffmann et al., 2020a), express ZAP and its cofactors (Figure S2A), and seemed most suitable for siRNA KD studies. Treatment with the different types of IFNs was associated with modest to marked increases in ZAP expression, and IFN-α and IFN-λ strongly induced ISG15 used as control of ISG stimulation (Figure 3A). Determination of virus yields by RT-qPCR showed that IFN-γ reduced virus production by almost 4 orders of magnitude at 100 U/ml (Figure 3B). IFN-β and IFN-λ were also highly potent against SARS-CoV-2, whereas IFN-α showed only modest inhibitory activity. Altogether, our data add to the recent evidence (Blanco-Melo et al., 2020; Mantlo et al., 2020) that IFNs are highly effective against SARS-CoV-2. However, they also revealed that at least in Calu-3 cells, type II IFN-γ is particularly effective and type I IFN-α only weakly active against SARS-CoV-2. In addition, our results show that ZAP and its cofactors are expressed in SARS-CoV-2 target cells and agree with a potential role of ZAP in the antiviral effect of the various IFNs.

**Figure 3:**
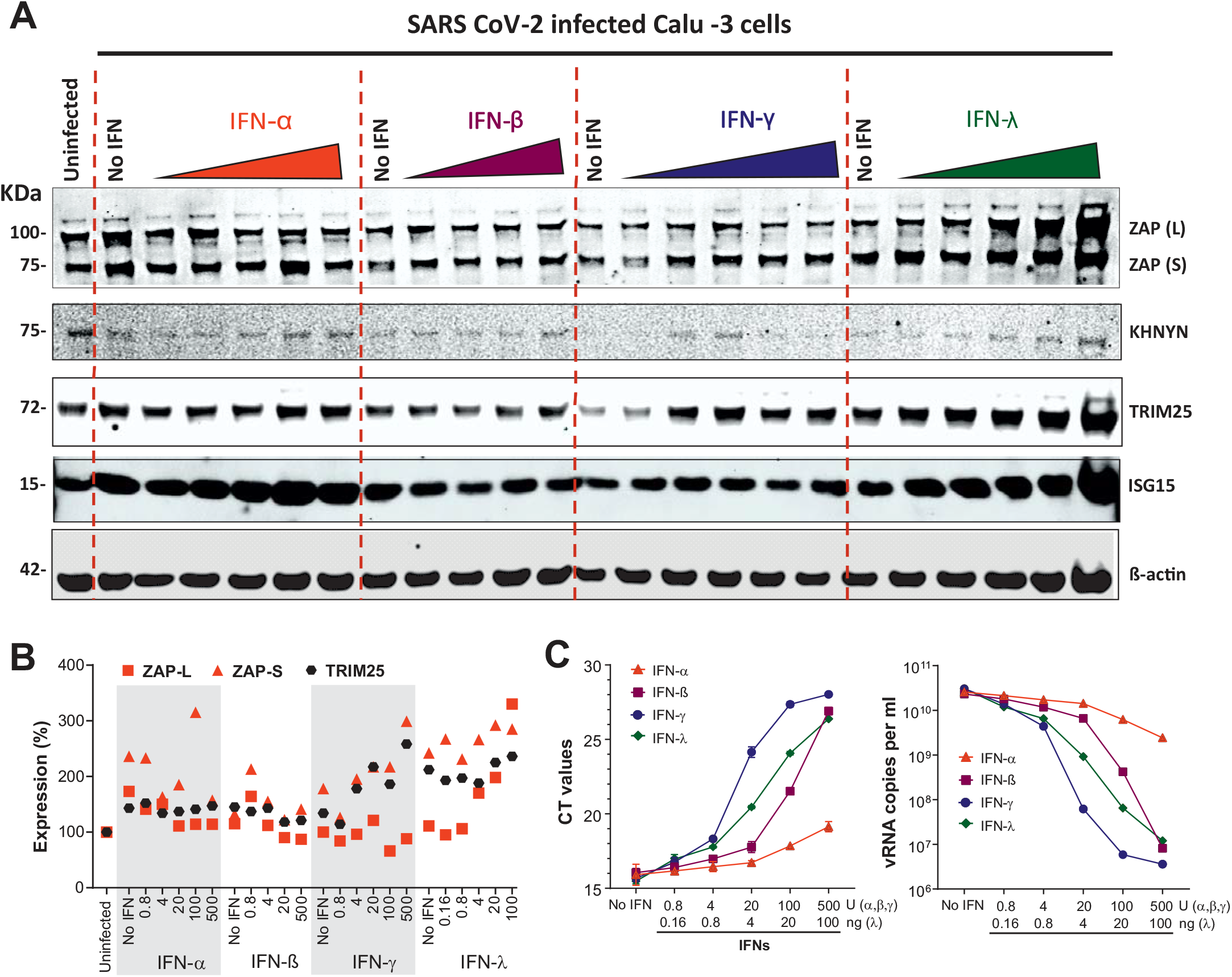
Inhibition of SARS-CoV-2 by different types of IFN. **(A)** Endogenous expression of ZAP and its cofactors KHNYN and TRIM25 in Calu-3 cells, that were left untreated and or uninfected or treated with the indicated amounts of IFNs in panel (B) and infected with SARS-CoV-2. Whole cell lysates were immunoblotted and stained with anti-ZAP, anti-KHNYN, anti-TRIM25, anti-ISG15 and anti-ß-actin. **(B)** Relative expression of the long (L) or short (S) isoforms of ZAP and TRIM25 normalized to unstimulated cells set as 100%. The data were derived from the Western blot shown in panel A. **(C)** Raw RT-qPCR CT values (left) and corresponding SARS-CoV-2 viral RNA copy numbers per mL (right) in the supernatants of Calu-3 cells as in (A). n= 1 in technical duplicates. See also Figure S2.

### Endogenous ZAP expression restricts SARS-CoV-2

To examine whether endogenous ZAP restricts SARS-CoV-2 and contributes to the antiviral effect of IFNs, we performed siRNA knock-down (KD) studies in Calu-3 cells and infected them. Western blot analyses showed that SARS-CoV-2 infection alone enhances expression of the short isoform of ZAP about 2-fold and this induction was further enhanced by IFN-ß and -γ treatment (Figure 4A, S3A, S3B). On average, treatment with ZAP siRNA reduced both ZAP(L) and ZAP(S) expression levels by ~60% without affecting TRIM25 and KHNYN expression levels (Figure 4A, S3A, S3B). In the initial experiment, siRNA-mediated KD of ZAP increased the levels of SARS-CoV-2 RNA determined by RT-qPCR (Figure S3C) in the absence of IFN by ~40% (Figure S3A). IFN-α treatment reduced virus yield ~337-fold and ZAP KD by ~80% increased viral RNA levels in the culture supernatants by 6.5-fold (Figure S3A). In agreement with the titration experiments (Figure 3B), IFN-β and IFN-γ were more effective than IFN-α and reduced SARS-CoV-2 production by ~4000-fold. IFN-λ was not available for the initial experiment and ZAP siRNA KD had no significant effect on virus yields upon treatment with IFN-β and IFN-γ.

**Figure 4:**
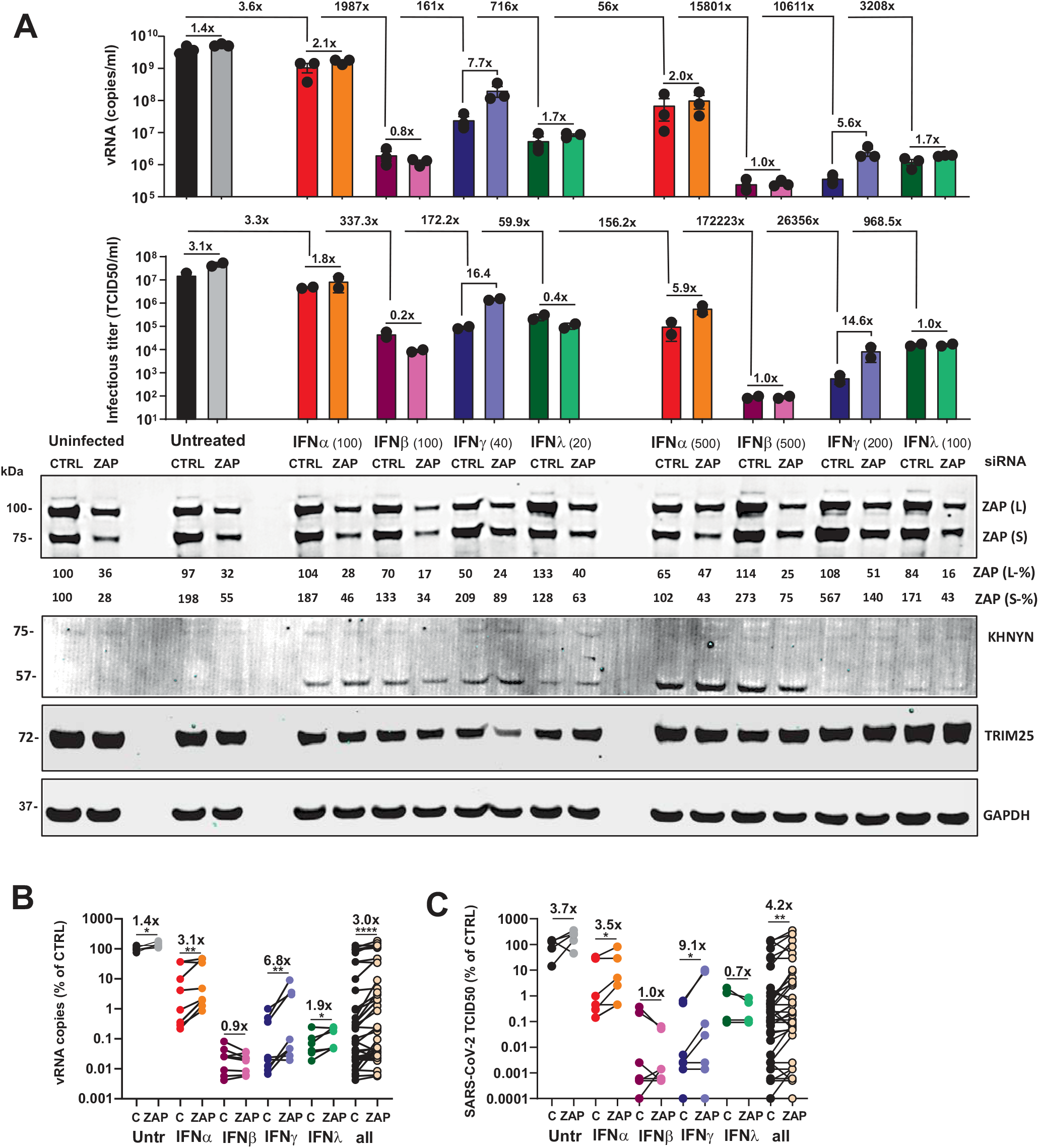
Role of ZAP in restricting SARS-CoV-2 production. **(A)** SARS-CoV-2 RNA levels (top panel, n= 3 (biological replicates) +/− SEM) and infectious titres (middle panel. n= 2 (biological replicates) +/− SD) in the supernatant of Calu-3 cells that were left untreated and/or uninfected or infected with SARS-CoV-2 and treated with the indicated amounts of IFNs. Cells were additionally transfected either with control or ZAP siRNA (CTRL and ZAP) as indicated. Immunoblots of whole cell lysates were stained with anti-ZAP, anti-KHNYN, anti-TRIM25 and anti-GAPDH (bottom panel). Relative expression of the long (L) or short (S) isoforms of ZAP are indicated below the blots. One representative blot of 2 biological replicates shown. **(B, C)** SARS-CoV-2 RNA copy numbers (B) or infectious virus titres (C), quantified relative to the control in the supernatant of Calu-3 cells that were left untreated or treated with the indicated IFNs and transfected with control (C) or ZAP (ZAP) siRNA. Numbers above the samples indicate the average change in vRNA or infectious titre. P-values, *, <0.05; **, <0.01; ***, <0.001; ****, <0.0001; unpaired Students t-test. See also Figure S1.

Saturating effects and almost complete inhibition of SARS-CoV-2 by other antiviral factors in the presence of IFN-β and IFN-γ might explain the lack of an effect of ZAP siRNA KD on virus yield. To further assess this, we repeated the ZAP siRNA KD experiment including 5-fold lower quantities of the different IFNs than in the previous setting and also included IFN-λ (Figure 4A). The results confirmed that IFN-β, -γ and -λ are substantially more effective against SARS-CoV-2 than IFN-α (Figure 4A). Again, ZAP KD slightly increased SARS-CoV-2 RNA levels in culture supernatants in the absence of IFNs and frequently more efficiently in its presence. The effects of ZAP KD upon IFNγ treatment were particularly pronounced (i.e. 7.7- and 5.6-fold) at the low and high dose, respectively (Figure 4A). On average under all conditions, ZAP KD increased SARS-CoV-2 RNA production by 3.0-fold (Figure 4B, right). The enhancing effect in the absence of IFN was modest (1.4-fold) but significant and consistent (Figure 4B, left). The effect of ZAP KD on vRNA yield was most pronounced in the presence of IFN-γ (6.8x) or IFN-α (3.1x) and modest (1.9x) or absent upon treatment with IFN-λ or IFN-β, respectively (Figure 4B). To further analyse the effects of IFN treatment and ZAP KD on SARS-CoV-2, we determined the infectious virus yields in the culture supernatants. Results of the TCID_50_ endpoint titration, although more variable, correlated well with the RT-qPCR data (Figure 4A, S3A; R^2^=0.713, p<0.0001). On average, ZAP KD increased infectious virus yield 4.2-fold. In agreement with the vRNA data, the enhancing effect was most pronounced in the presence of IFN-γ (9.1x) and absent upon treatment with IFN-β or IFN-λ (Figure 4C). The effects of ZAP KD on SARS-CoV-2 RNA yield and infectious titres were most obvious at non-saturating levels of IFNs (Figure 4B, 4C). Altogether, the results clearly demonstrated that endogenous ZAP restricts SARS-CoV-2, especially in the presence of IFN-γ.

## DISCUSSION

Coronaviruses generally show CpG suppression and in agreement with recent findings (Xia, 2020), we found that the SARS-CoV-2 genome is particularly poor in CpG dinucleotides. Remarkably, the closest bat relatives of SARS-CoV-2 show the strongest suppression and lowest frequency of CpGs among all available bat CoV genomes. Thus, zoonotic transmission of a bat or intermediate host CoV strain with unusually low CpG content may have facilitated the effective spread of SARS-CoV-2 in humans. On average, CpGs are more strongly suppressed in humans than in bats suggesting that the selection pressure for CpG suppression might be increased after zoonotic transmission. In agreement with this possibility, we show that the antiviral factor ZAP that specifically targets CpG dinucleotides is expressed in human viral target cells and restricts SARS-CoV-2. Knock-down of ZAP in viral target cells moderately enhanced SARS-CoV-2 production in the absence of IFN but had much stronger effects in the presence of type II IFN-γ that was highly active against this viral pathogen. Altogether, our data clearly show that ZAP is one of the cellular effectors that contribute to the strong anti-SARS-CoV-2 activity of IFNs.

Consistent with an increased selection pressure for CpG suppression in the human host, we found that community-acquired hCoVs show lower frequencies of CpG dinucleotides than their closest animal relatives (Figure 2A). This was not observed for highly pathogenic SARS- and MERS-CoVs most likely reflecting less advanced human adaptation consistent with their less effective and transient spread. In addition, selection pressures may not act on all parts of the genome. Specifically, while SARS-CoV-2 shows low CpG frequencies throughout most parts of its genome the number of CpGs at the 3`end is high. Notably, several ORFs overlap in this part of the genome, which might make it difficult for the virus to get rid of CpGs without fitness cost. However, this may render SARS-CoV-2 vulnerable to ZAP restriction since many coronavirus mRNA transcripts encompass this region (Kim et al., 2020). Interestingly, it has been reported that almost all changes in the N gene, which contains higher numbers of CpGs (Figure 2D), emerging during spread of SARS-CoV-2 in the human population eliminate these dinucleotides, while this is less common in other parts of the genome (Gioacchino et al., 2020). Recent data showed that a region of CpGs at the beginning of the *env* gene of HIV-1 rather than overall genomic content determine the susceptibility of HIV-1 to inhibition by ZAP (Kmiec et al., 2020). Thus, it is conceivable that parts of the SARS-CoV-2 genome may still be ZAP sensitive, although it shows strong CpG suppression throughout most parts of its genome. Further studies on the evolutionary constrains acting on SARS-CoV-2 and further elimination of CpGs during human adaptation will be interesting. They might also reveal whether selection pressure for loss of CpGs may promote the emergence of less virulent virus variants. For example, SARS-CoV-2 contains a unique potential furin cleavage site in its Spike protein that is absent in bat and pangolin viruses and introduces several CpG dinucleotides (Figure 2F). Increased furin-mediated activation of the Spike protein might affect viral infectivity as well as cell tropism and consequently its pathogenicity (Coutard et al., 2020; Hoffmann et al., 2020b; Tse et al., 2014). It has been observed that this site may acquire mutations during viral passage in cell culture (Ogando et al., 2020) and it will be of interest to determine whether this insertion increases ZAP sensitivity and might also be prone to mutation or elimination *in vivo* (Andres et al. 2020).

SARS-CoV-2 is most closely related to two bat viruses (RaTG13 and RmYN02), which show about 96% sequence identity to the human virus (Zhou et al., 2020a, 2020b). However, the degree of sequence homology is unevenly distributed throughout the viral genomes and it is under debate whether SARS-CoV-2 might represent a recombination between CoVs found in the reservoir bat host and viruses found in intermediate hosts, such as pangolins, that also show high sequence identity to the human virus (Gu et al., 2020; Lam et al., 2020). SARS-CoV-2 differs significantly from its bat relatives in the Spike coding region. Notably, we found that the RmYN02 bat CoV strain that is otherwise highly related to SARS-CoV-2 shows substantially higher CpG frequencies in the region encoding the ACE2 receptor binding region of the viral Spike (Figure 2E). In this part of the genome SARS-CoV-2 shows substantially higher similarity in sequence and CpG numbers to bat RaTG13 and pangolin Pang Guangdong CoV strains than to RmYN02. These differences need further investigation and the possibility that recombination may have facilitated the loss of regions with high CpG frequencies warrants further investigation.

In agreement with recent data (Mantlo et al., 2020), we found that IFN-β efficiently inhibits SARS-CoV-2, while IFN-α was less effective. However, IFN-λ was about as effective as IFN-β. Most notably and in agreement with recent studies in colon organoids (Stanifer et al., 2020), type III IFN-γ showed the highest efficacy against SARS-CoV-2 (Figure 3). More studies on the anti-SARS-CoV-2 activity of the various types of IFNs and their potential adverse effects in patients are required to optimize IFN-based therapeutic approaches. At least in Calu-3 cells IFN-γ displayed the highest potency against SARS-CoV-2. IFN-γ has been used in a wide variety of clinical indications and a tendency for higher levels of IFN-γ in moderate compared to severe cases of COVID-19 has been reported (Chen et al., 2020). Thus, further studies on the application of IFN-γ in the treatment of COVID-19 are highly warranted.

Both SARS-CoV-2 infection alone as well as IFN treatment induced ZAP expression in our cell-based systems. This result agrees with the recent finding that ZAP mRNA expression is clearly (i.e. 8-fold) induced in SARS-CoV-2-infected human individuals (Blanco-Melo et al., 2020). In agreement with previous studies (Li et al., 2019), especially the short isoform of ZAP was induced by virus infection and IFN treatment, whereas the long isoform was constitutively expressed at relatively high levels but hardly inducible. Knock-down of ZAP by ~60% increased SARS-CoV-2 RNA yield by ~40% in the absence of IFN. The enhancing effect of ZAP KD on SARS-CoV-2 RNA yield and infectious titres varied to some extent, at least in part due to variations in KD efficiencies and saturating effects in the presence of high of IFN-β, -γ and -λ. In addition, the type of IFN seems to play a significant role. On average, ZAP KD increased SARS-CoV-2 RNA and infectious virus yield by 6.8- and 9.1-fold upon treatment with IFN-γ but had no significant enhancing effect in the presence of IFN-β. In part, this can be explained by the fact that IFN-γ induced ZAP(S) expression in Calu-3 with higher efficiency than other IFNs. It will also be interesting to clarify whether IFN-β might be particularly effective in inducing other antiviral factors masking effects of ZAP. IFN treatment induces hundreds of potentially antiviral ISGs. Thus, it is remarkable that on average ZAP KD increased SARS-CoV-2 RNA yield and infectious titres ~6-fold in the presence of IFN levels resulting in robust but non-saturating (i.e. 10- to 500-fold) inhibition of SARS-CoV-2. These results clearly show that ZAP contributes to the antiviral effect of IFNs. They further support a significant role of the short isoform of ZAP in restricting SARS-CoV-2 because it is more responsive to IFN treatment than the long isoform. It has been reported that mainly the long isoform of ZAP exerts antiviral activity and that the short isoform even down-tunes the antiviral IFN response in a negative-feedback mechanism (Schwerk et al., 2019). However, the roles of the different isoforms of ZAP in innate antiviral immunity are under debate and their effects on SARS-CoV-2 clearly need to be addressed in future studies.

It has been reported that ZAP itself lacks RNase activity and requires cofactors, i.e. TRIM25 and KHNYN, for effective restriction of RNA viruses (Ficarelli et al., 2019; Li et al., 2017; Zheng et al., 2017). Both of these cellular factors were detected in the lung cells used in the present study and not affected by ZAP siRNAs. However, the detectable levels of KHNYN were relatively low and it seems that a significant proportion of it might be cleaved. Further investigation is necessary but the results are somewhat reminiscent of MALT-4-mediated cleavage of the antiretroviral RNAse N4BP1 in activated T cells (Yamasoba et al., 2019). Despite low detectable levels of full-length KHNYN, however, knock-down of ZAP clearly enhanced SARS-CoV-2 RNA yield. Thus, either the residual levels are sufficient for anti-SARS-CoV-2 activity or related RNases, may serve as ZAP cofactors in virus restriction (Nchioua et al., 2020).

In summary, our results show that ZAP restricts SARS-CoV-2 and contributes to its inhibition by IFNs. They further suggest that zoonotic transmission of a CoV already showing strong CpG suppression may have facilitated effective spread in humans. Our data also confirm that SARS-CoV-2 is highly susceptible to IFNs and might motivate assessment of combination therapies including IFN-γ for treatment of COVID-19. If SARS-CoV-2 is highly sensitive to IFN, why is it frequently not efficiently controlled by the innate immune response? It is known that coronaviruses use various mechanisms to avoid immune sensing (Fehr and Perlman, 2015; Park and Iwasaki, 2020; Totura and Baric, 2012) and recent data show that the SARS-CoV-2 Nsp1 proteins blocks ribosomal translation of cellular proteins including antiviral defence factors (Thoms et al., 2020). However, we are still far from a full understanding of viral immune evasion and counteraction mechanisms. ZAP is only one of numerous effectors of the antiviral immune response. Many others remain to be defined and it will be important to determine whether the different types of IFN use different effectors to restrict SARS-CoV-2. A better knowledge of the anti-SARS-CoV-2 effectors of the IFN response and the viral countermeasures will help to develop safe and effective immune therapy approaches against COVID-19.

## AUTHOR CONTRIBUTIONS

R.N. performed most experiments and D.K. performed phylogenetic and CpG frequency analyses. J.M. and C.C established and performed experiments with infectious SARS-CoV-2. R.G. determined viral RNA copy numbers. C.S., S.N. and S.S. provided resources. D.S., J.M., and K.M.J.S. helped with the design of the experiments and interpretation of the results. F.K., R.N. and D.K. designed the experiments and wrote the manuscript. All authors reviewed and proofread the manuscript.

## ACKNOWLEDGMENTS

We thank Susanne Engelhart, Kerstin Regensburger, Martha Meyer, Regina Burger, Nicola Schrott and Daniela Krnavek for excellent technical assistance. This study was supported by DFG grants to F.K., J.M., D.K., D.S., S.S. and K.S. (CRC 1279, SPP 1923, KM 5/1-1, SP1600/4-1), EU’s Horizon 2020 research and innovation program to J.M. (Fight-nCoV, 101003555), as well as intramural funding by the Ulm University Medical Center (L.SBN.0150) to K.S. C.C., and R.G. are part of and R.G. is funded by a scholarship from the International Graduate School in Molecular Medicine Ulm.

## STAR METHODS

### CONTACT FOR REAGENT AND RESOURCE SHARING

#### Lead contact

Further information and requests for resources and reagents should be directed to and will be fulfilled by the Lead Contact, Frank Kirchhoff.

#### Materials Availability

All unique reagents generated in this study are listed in the Key Resources Table and available from the Lead Contact.

## EXPERIMENTAL MODEL AND SUBJECT DETAILS

### Ethical statement for human samples

No patient-derived primary cells were used in the present study. The use of established cell lines did not require the approval of the Institutional Review Board.

### Phylogenetic analyses

Nucleotide sequences of full-length or near full-length coronavirus genomes were obtained from the NCBI or GISAID (Koehorst J., van Dam JCJ., Saccenti E., Martins dos Santos V., 2017) database and aligned using Clustal Omega (https://www.ebi.ac.uk/Tools/msa/clustalo/). Phylogenetic trees showing distance-based relationship inference based on representative full genome nucleotide sequences were generated using NGPhylogeny.fr FastME tool (https://ngphylogeny.fr/) (Lemoine et al., 2019), and visualized using iTOL (https://itol.embl.de/).

### Cell culture

Vero E6 (*Cercopithecus aethiops* derived epithelial kidney cells, ATCC) cells were grown in Dulbecco’s modified Eagle’s medium (DMEM, Gibco Cat#41965039) which was supplemented with 2.5% heat-inactivated fetal calf serum (FCS, Gibco Cat#10270106), 100 units/ml penicillin, 100 μg/ml streptomycin (ThermoFisher, Cat#15140122), 2 mM L-glutamine, 1 mM sodium pyruvate (Pan Biotech, Cat# P04-8010), and 1x non-essential amino acids (Sigma, Cat#M7145). Caco-2 (human epithelial colorectal adenocarcinoma, kindly provided by Prof. Barth, Ulm University) cells were grown in the same media but with supplementation of 10% FCS. Calu-3 (human epithelial lung adenocarcinoma, kindly provided by Prof. Frick, Ulm University) cells were cultured in Minimum Essential Medium Eagle (MEM, Sigma Cat#M4655) supplemented with 10% FCS (during viral infection) or 20% (during all other times), 100 units/ml penicillin, 100 μg/ml streptomycin, 1 mM sodium pyruvate, and 1x non-essential amino acids. NHLF (primary human lung fibroblasts, Lonza) cells, and A549 (adenocarcinomic human alveolar basal epithelial cells, ATCC) were cultured in DMEM supplemented with 10% FCS, 2 mM μg/ml L-glutamine, 100 units/ml penicillin and 100 μg/ml streptomycin.

### CpG content sliding window analysis

Representative full genomic sequences of Coronaviruses (NCBI accession numbers listed in Table 1) were aligned using DECIPHER, with a gap opening/extension penalty of 70, to avoid formation of large gaps. Numbers of CpGs were extracted using a sliding window of 100 nucleotides and a step size of 100. All analyses were done using R. The raw data is displayed as a heat map generated by Graphpad PRISM.

### Species specific CpG content

mRNA sequences were extracted from genome transcripts (NCBI) of Homo sapiens (assembly GRCh38.p13), Rhinolophus ferrumequinum (assembly mRhiFer1_v1.p), Manis javanica (assembly ManJav1.0), Camelus dromedarius (assembly CamDro3), Bos taurus (assembly ARS-UCD1.2), Sus scrofa (assembly Sscrofa11.1), Rattus rattus (assembly Rrattus_CSIRO_v1). CpG suppression was calculated as [number of CpG * mRNA sequence length]/[number of C * number of G].

### qRT-PCR

*N* (nucleoprotein) transcript levels were determined in supernatants collected from SARS-CoV-2 infected Calu-3 cells 48 h post-infection. Total RNA was isolated using the Viral RNA Mini Kit (Qiagen, Cat# 52906) according to the manufacturer’s instructions. RNA concentrations were determined using the NanoDrop 2000 Spectrophotometer. RT-qPCR was performed as previously described (Groß et al., 2020) using TaqMan® Fast Virus 1-Step Master Mix (Thermo Fisher, Cat#4444436) and a OneStepPlus Real-Time PCR System (96-well format, fast mode). Primers were purchased from Biomers (Ulm, Germany) and dissolved in RNAse free water. Synthetic SARS-CoV-2-RNA (Twist Bioscience, Cat#102024) or RNA isolated from BetaCoV/France/IDF0372/2020 viral stocks quantified via this synthetic RNA (for low Ct samples) were used as a quantitative standard to obtain viral copy numbers. All reactions were run in duplicates. (Forward primer (HKU-NF): 5’-TAA TCA GAC AAG GAA CTG ATT A-3’; Reverse primer (HKU-NR): 5’-CGA AGG TGT GAC TTC CAT G-3’; Probe (HKU-NP): 5’-FAM-GCA AAT TGT GCA ATT TGC GG-TAMRA-3’.

### SDS-PAGE and Western blotting

SDS-PAGE and Western Blotting were performed as previously described (Kmiec et al.,2020). In brief, two days post-infection, cells were washed with PBS and lysed with Co-IP buffer. The samples were separated on 4-12% Bis-Tris gradient acrylamide gels (Invitrogen) and blotted onto polyvinylidene difluoride (PVDF). Blotted membranes were probed with anti-ZAP (GeneTex, Cat#GTX120134;) diluted 1 to 1000, anti-KHNYN (Santa Cruz Biotechnology, Cat#sc-514168) diluted 1 to 50, anti-TRIM25 (BD Biosciences, Cat# 610570) and anti-GAPDH (Biolegend, Cat# 607902) or anti-beta Actin (abcam, Cat#ab8227) both diluted 1 to 2000. Subsequently, blots were stained with IRDye® 680RD Goat anti-Rabbit IgG (H + L) (LI-COR, Cat #926-68071), IRDye® 800CW Goat anti-Mouse IgG (H + L) and IRDye® 800CW Goat anti-RAT IgG (H + L) (LI-COR, Cat #925-32219) secondary antibodies, all diluted 1 to 20.000, and scanned using a LI-COR Odyssey reader.

### Effect of IFNs on SARS-CoV-2 replication

300.000 Calu-3 cells were seeded in 12-well plates. 24h and 96h post-seeding cells were stimulated with increasing amounts of IFNs (α2, β and γ, 0.8, 4, 20, 100 and 500 U/ml) or IFN-λ-1 (0.16, 0.8, 4, 20 and 100ng) per 1 ml of MEM medium. 24 h post the first stimulation, the medium was exchanged. 2 h after the second stimulation Calu-3 cells were infected with SARS-CoV-2 (MOI 0.05) and 7 to 9 h later, supernatant was removed and 1 ml fresh medium was added.48h post-infection cells were harvested for further analysis.

### ZAP knockdown and IFN treatment in Calu-3 cells

300.000 Calu-3 cells were seeded in 12-well plates. 24 h and 96 h post-seeding they were transfected with either a non-targeting control (Eurofins, UUC UCC GAA CGU GUC ACG UdT dT) siRNA or ZAP-specific siRNA (siRNA SMART pool, Dharmacon, Cat#SO-2863397G) (Takata et al., 2017). 20μM siRNA was transfected in one well using Lipofectamine RNAiMAX (Thermo Fisher) according to the manufacturer’s instructions. Prior to transfection, the medium was changed. 14 h post transfection, the medium was replaced with 1 ml MEM supplemented with IFNs (100 or 500 U/ml IFN-α2, 100 or 500 U/ml IFN-β, 40 or 200 U/ml IFN-γ and 20 or 100ng of IFN-λ1). 7 h post transfection, the Calu-3 cells were infected with SARS-CoV-2 (MOI 0.05) and 7 to 9 h later, supernatant was removed and 1 ml fresh medium was added. 48h post infection cells and supernatants were harvested for further analysis.

### Virus strains and propagation

BetaCoV/Netherlands/01/NL/2020 was obtained from the European Virus Archive. The virus was propagated by infecting 70% confluent Vero E6 in 75 cm^2^ cell culture flasks at an MOI of 0.003 in 3.5 ml serum-free medium containing 1 μg/ml trypsin. The cells were incubated for 2 h at 37°C, before adding 20 ml medium containing 15 mM HEPES. The DMEM medium was changed after 3 days post infection and the supernatant harvested 48 h post infection upon visible cytopathic effect (CPE). Supernatants were centrifuged for 5 min at 1,000 × g to remove debris, aliquoted and stored at −80°C. The infectious virus titre was determined as plaque forming units (PFU) or TCID50. Genomic RNA copies were determined by RT-qPCR.

### Plaque-forming Unit (PFU) assay

To determine plaque forming units (PFU), SARS-CoV-2 stocks were serially diluted 10-fold. Monolayers of Vero E6 cells in 12-wells were infected with the dilutions and incubated for 1 to 3 h at 37°C with shaking every 15 to 30 min. Afterwards, the cells were overlayed with 1.5 ml of 0.8% Avicel RC-581 (FMC Corporation) in medium and incubated for 3 days. Cells were fixed by adding 1 ml 8% paraformaldehyde (PFA) and incubated at room temperature for 45 min. After discarding the supernatant, the cells were washed with PBS once, and 0.5 ml of staining solution (0.5% crystal violet and 0.1% triton in water) was added. After 20 min incubation at room temperature, the staining solution was removed using water, virus-induced plaque formation quantified, and PFU per ml calculated.

### TCID50 endpoint titration

SARS-CoV-2 stocks or infectious supernatants were serially diluted. 25,000 Caco-2 cells were seeded per well in 96 flat bottom well plates in 100 μl medium and incubated over night before 62 μl fresh medium was added. Next, 18 μl of titrated SARS-CoV-2 stocks or supernatant were used for infection, resulting in final dilutions of 1:10^1^ to 1:10^9^ on the cells in triplicates. Cells were then incubated for 6 days and monitored for CPE. TCID_50_/ml was calculated according to Reed and Muench method (REED and MUENCH, 1938).

### Calu-3, A549 and NHLF stimulation

Calu-3, A549 and NHLF cells were seeded in 12 well plates and stimulated with IL-2 (10ng/ml), PHA (25ng/ml), IL-27 (5ng/ml), TNFα (25ng/ml), IFN-α2 (500u/ml), IFN-β (500u/ml), IFN-γ (200u/ml) and IFN-λ1 (100ng). 3- and 6-days post stimulation cells were harvested and lysed for Western blot analysis. Calu-3 cells were stimulated for 24h only due to cytopathic effects of the IFN treatment.

## QUANTIFICATION AND STATISTICAL ANALYSIS

Statistical analyses were performed with GraphPad PRISM (GraphPad Software) and Microsoft Excel. P values were calculated using the two-tailed unpaired Student’s-t-test unless specified otherwise. Correlations were calculated with the linear regression module. Unless otherwise stated, all experiments were performed in triplicate and the data are shown as mean ± SEM. Significant differences are indicated as: *p < 0.05; **p < 0.01; ***p < 0.001. Statistical parameters are specified in the figure legends.

## KEY RESOURCES TABLE

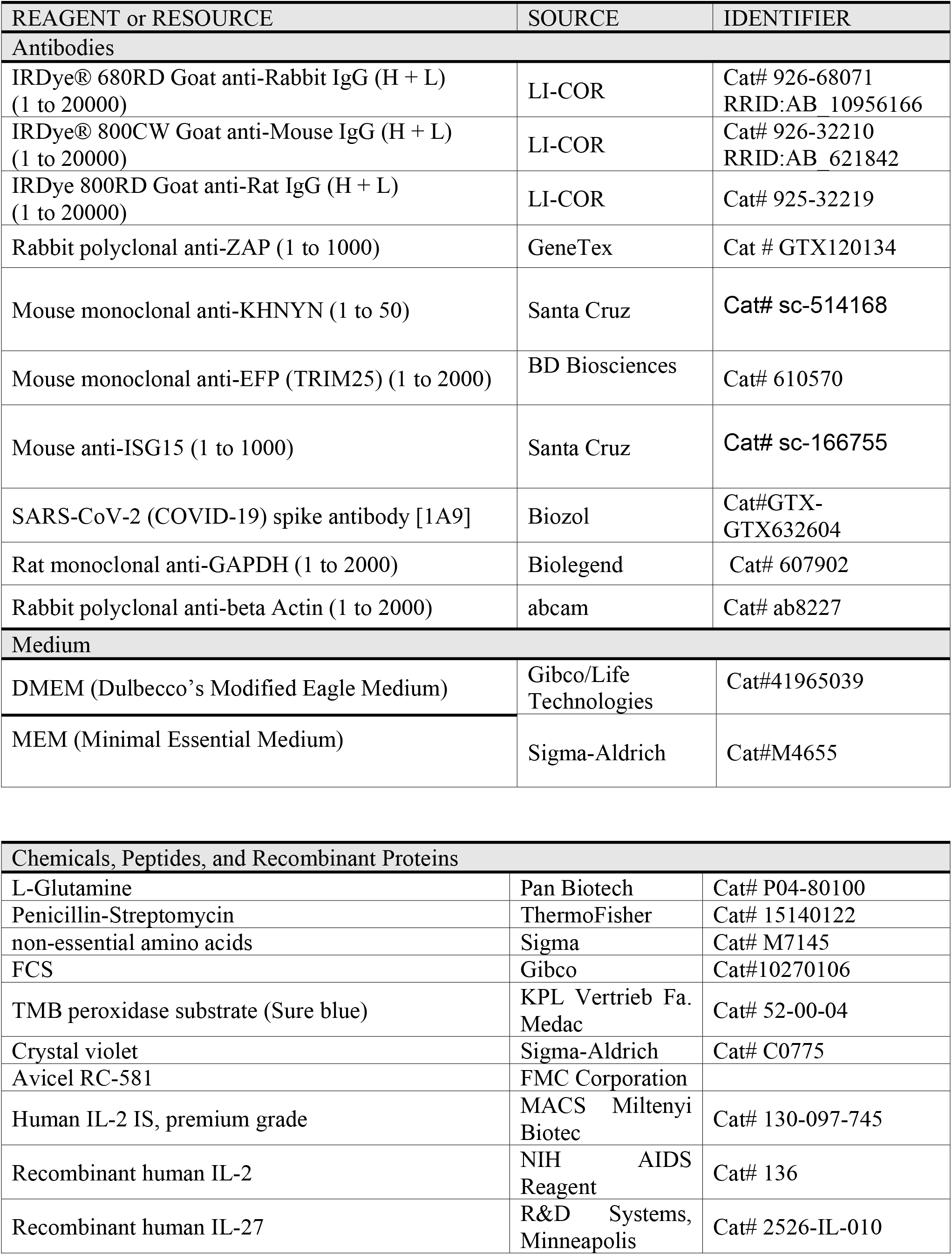
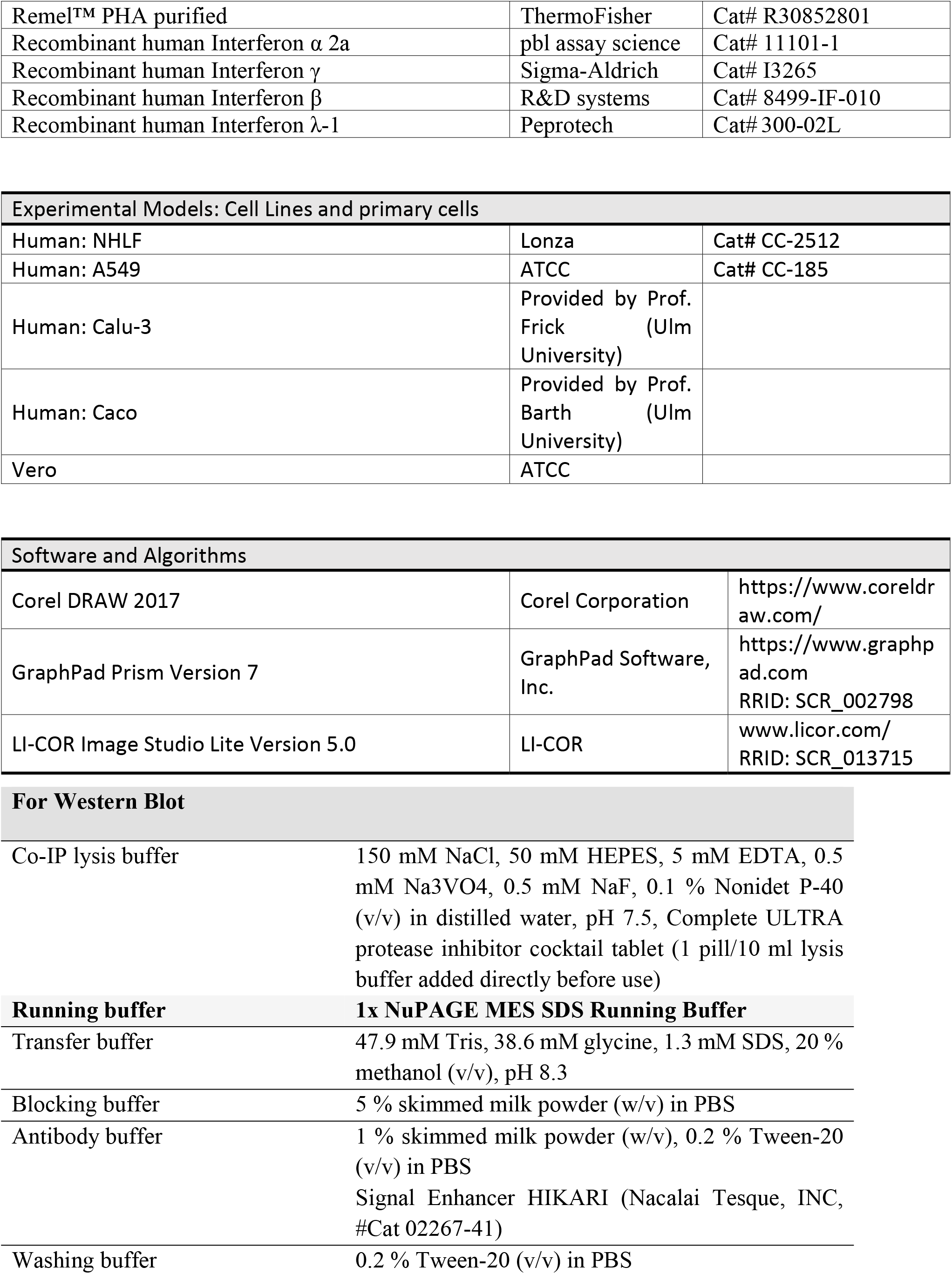

## SUPPLEMENTARY INFORMATION

**Figure S1:**
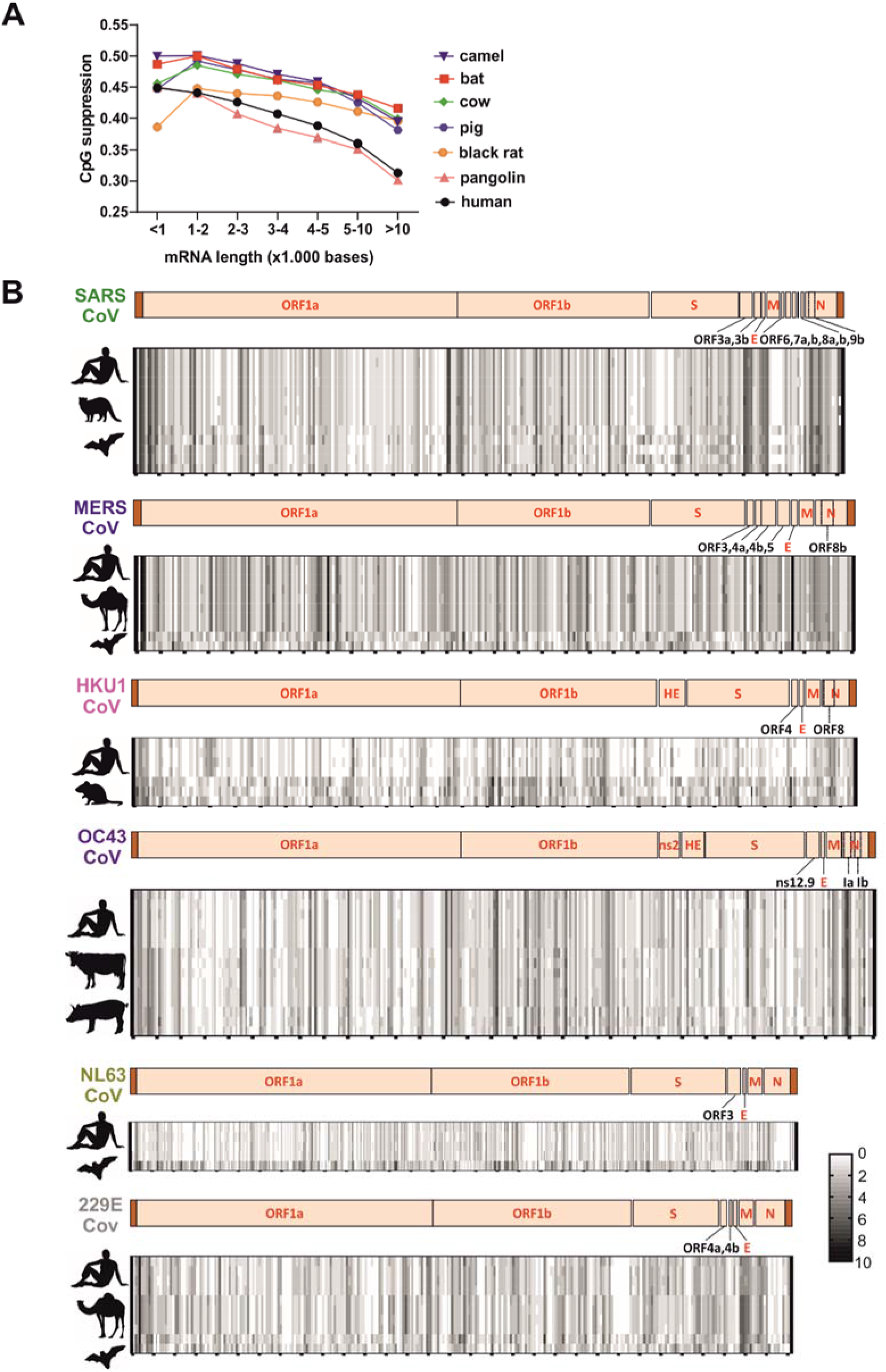
Length-dependent CpG suppression in host mRNAs and CpG distribution in the genomes of human and animal coronaviruses (related to Figure 2). **(A)** CpG suppression in host mRNA transcripts grouped by average length as indicated. **(B)** CpG frequency to human and animal Coronavirus genomes. The colour shade indicates the amount of CpGs per 100 nucleotides (0 (white) to 10 (black)). Genome organisation diagrams above represent the reference human virus as indicated.

**Figure S2:**
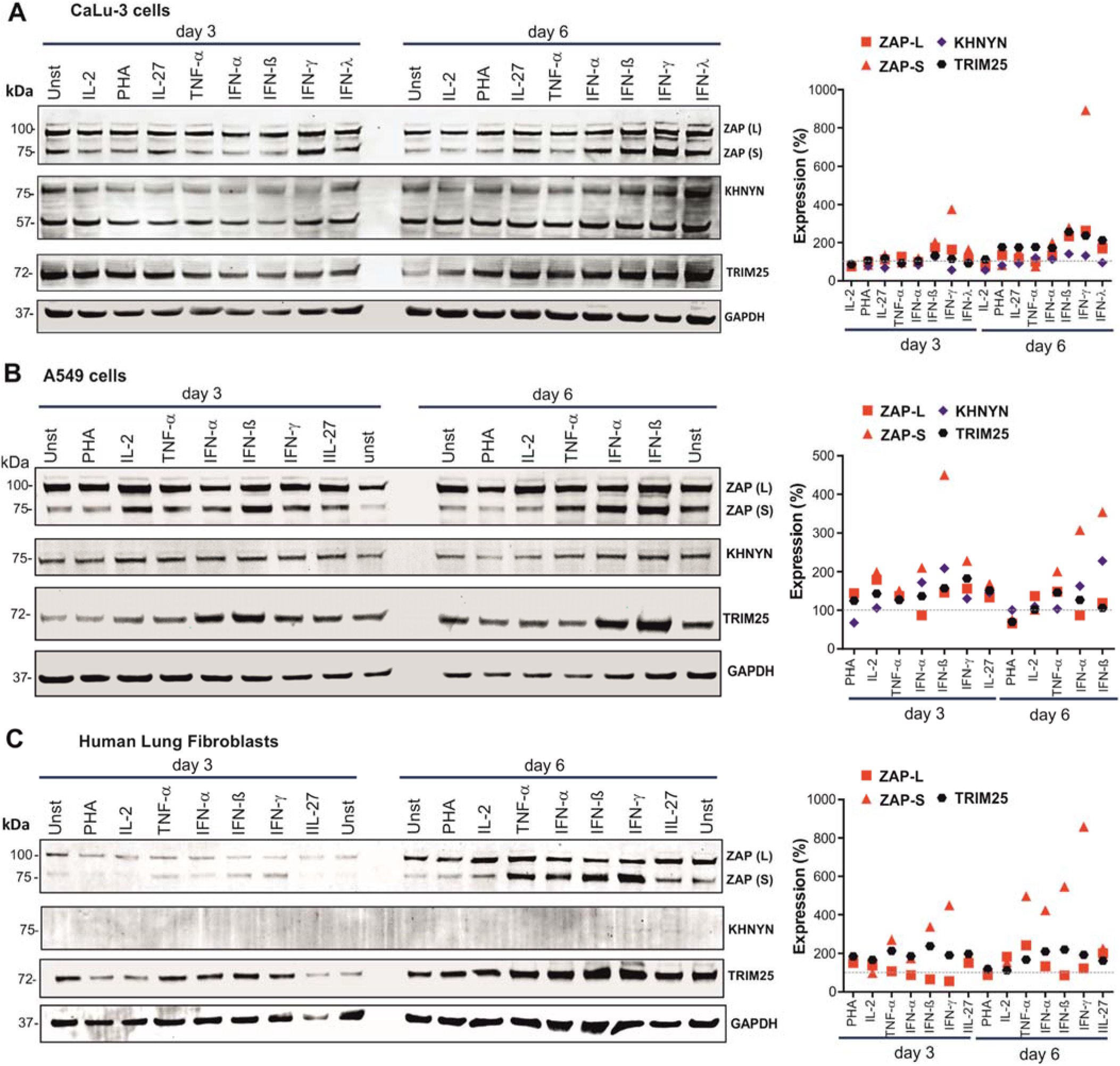
Expression and IFN induction of ZAP and its cofactors KHNYN and TRIM25 (related to Figure 3). **(A-C)** Expression of endogenous ZAP, KHNYN and TRIM25 in (A) Calu-3 (B) A549 C) primary human lung fibroblasts (NHLF), and indicated cytokines. Immunoblot of whole cell lysates stained with anti-ZAP, anti-KHNYN, anti-TRIM25. GAPDH serves as a protein loading control. Relative expression of the long (L) or short (S) isoforms of ZAP, KHNYN and TRIM25 normalized to unstimulated cells set as 100%, are indicated in the right panels. For NHLF KHNYN was not quantified because the levels were to for definitive quantification.

**Figure S3:**
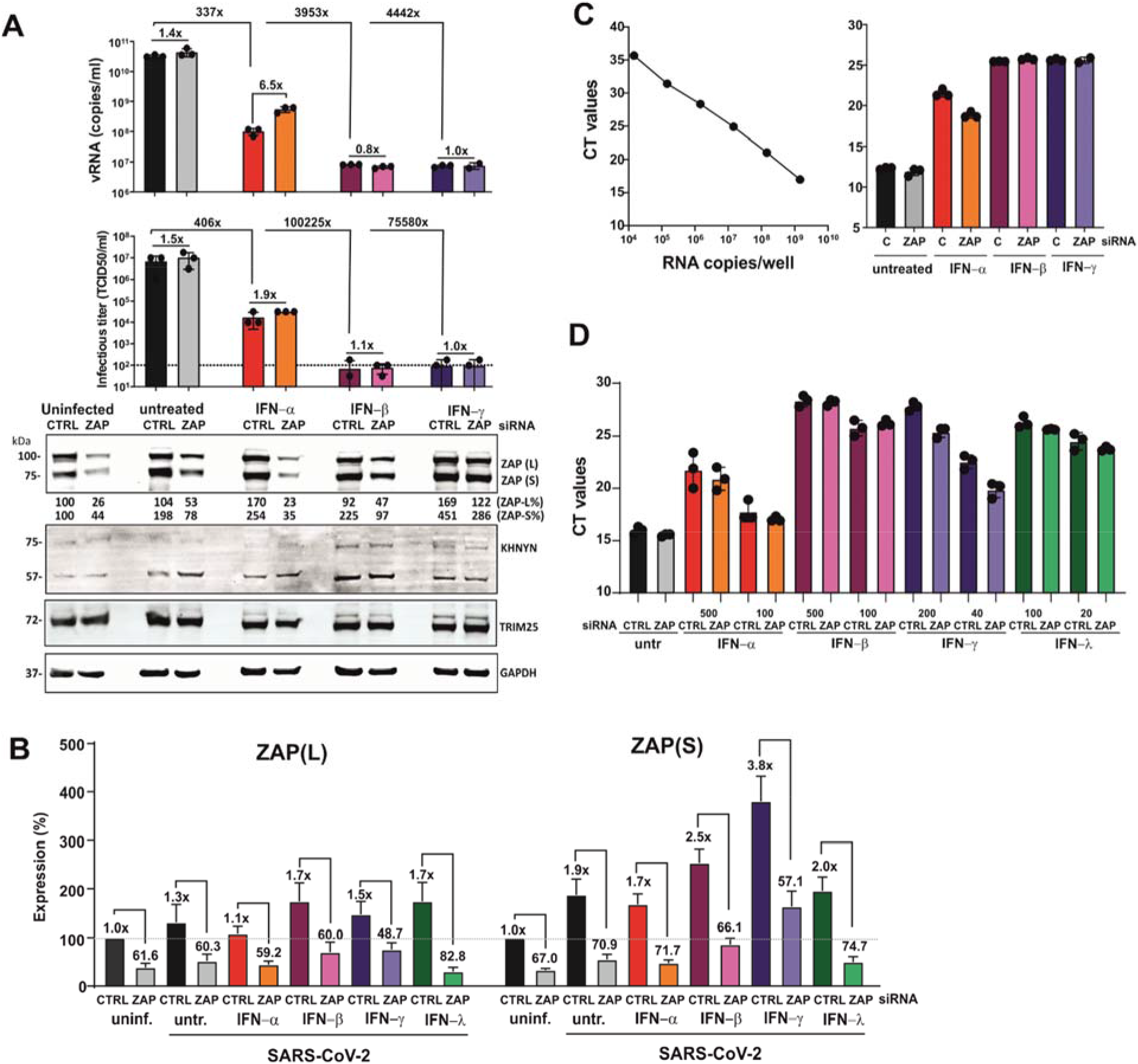
Expression and anti-SARS-CoV-2 activity of ZAP (related to Figure 4). (**A**) SARS-CoV-2 RNA levels (top panel, n= 3 (biological replicates) +/− SD) and infectious titres (middle panel. n= 3 (biological replicates) +/− SD) in the supernatant of Calu-3 cells that were left untreated and/or uninfected or infected with SARS-CoV-2 and treated with the indicated amounts of IFNs. Cells were additionally transfected either with control or ZAP siRNA (CTRL and ZAP) as indicated. Immunoblots of whole cell lysates were stained with anti-ZAP, anti-KHNYN, anti-TRIM25 and anti-GAPDH (bottom panel). Relative expression of the long (L) or short (S) isoforms of ZAP are indicated below the blots. One representative blot of 2 biological replicates shown. (**B**) ZAP(L) and ZAP(S) expression levels in Calu-3 cells upon infection and IFN treatment relative to expression levels in untreated cells (100%). (**C**) Standard curve and raw qRT-PCR CT values corresponding to the SARS-CoV-2 RNA copy numbers per mL shown in panel A. (**D**) Raw qRT-PCR CT values for the viral RNA levels shown in Figure 4A. Panels B and C show mean values (+/− SD) from three replicates.

**Supplemental Table 1.** Features of Coronavirus sequences analyzed (related to Figure 1).

| **Viral strain** | **seq length** | **no of CpG** | **CpG frequency** | **CpG suppression** | **GC content %** | **% identity to ref.** | **accession no./ID** |
| --- | --- | --- | --- | --- | --- | --- | --- |
| **SARS-hCoV-2 Wuhan-Hu-1** | **29872** | **436** | **0,0146** | **0,406** | **38,0** |  | **NC 045512.2** |
| SARS-hCoV-2 TWN/CGMH-CGU-01/2020 | 29862 | 439 | 0,0147 | 0,408 | 38,0 | 99 | MT192759.1 |
| SARS-hCoV-2 USA/MN3-MDH3/2020 | 29783 | 439 | 0,0147 | 0,409 | 38,0 | 99 | MT188339.1 |
| SARS-hCoV-2 BRA/SP02cc/2020 | 29903 | 440 | 0,0147 | 0,409 | 38,0 | 99 | MT350282.1 |
| bat CoV RaTG13 | 29855 | 441 | 0,0148 | 0,409 | 38,0 | 96 | MN996532.1 |
| bat SARS RmYN02 | 29671 | 451 | 0,0152 | 0,416 | 38,2 | 93 | EPI ISL 412977 |
| pangolin CoV PCoV GX-P2V | 29795 | 461 | 0,0155 | 0,418 | 38,5 | 85 | MT072864.1 |
| pangolin CoV PCoV GX-P1E | 29801 | 459 | 0,0154 | 0,416 | 38,5 | 85 | MT040334.1 |
| pangolin CoV-19/pangolin/Guangdong/1/2019 | 29825 | 428 | 0,0144 | 0,393 | 38,2 | 90 | EPI ISL 410721 |
| **SARS hCoV Tor2** | **29751** | **568** | **0,0191** | **0,460** | **40,8** |  | **NC 004718.3** |
| SARS hCoV GD01 | 29757 | 570 | 0,0192 | 0,460 | 40,8 | 99 | AY278489.2 |
| SARS hCoV CUHK-W1 | 29736 | 567 | 0,0191 | 0,459 | 40,8 | 99 | AY278554.2 |
| SARS hCoV ZS-C | 29647 | 566 | 0,0191 | 0,460 | 40,8 | 99 | AY395003.1 |
| civet SARS CoV SZ16 | 29731 | 569 | 0,0191 | 0,459 | 40,8 | 99 | AY304488.1 |
| civet SARS CoV civet007 | 29540 | 569 | 0,0193 | 0,462 | 40,8 | 99 | AY572034.1 |
| civet SARS CoV SZ3 | 29741 | 569 | 0,0191 | 0,459 | 40,8 | 99 | AY304486.1 |
| civet SARS CoV civet010 | 29518 | 568 | 0,0192 | 0,461 | 40,8 | 99 | AY572035.1 |
| bat CoV WIV16 | 30290 | 596 | 0,0197 | 0,471 | 40,9 | 94 | KT444582.1 |
| bat CoV BtRs-BetaCoV/YN2018A | 29698 | 596 | 0,0201 | 0,477 | 41,0 | 93 | MK211375.1 |
| bat CoV BtRs-BetaCoV/YN2018B | 30256 | 588 | 0,0194 | 0,466 | 40,8 | 94 | MK211376.1 |
| bat CoV BtRs-BetaCoV/YN2018C | 29689 | 589 | 0,0198 | 0,470 | 41,1 | 94 | MK211377.1 |
| bat CoV Rs4874 | 30311 | 597 | 0,0197 | 0,471 | 40,9 | 94 | KY417150.1 |
| **MERS hCoV-EMC** | **30119** | **711** | **0,0236** | **0,555** | **41,2** |  | **NC 019843.3** |
| MERS hCoV Al-Hasa 1 2013 | 30117 | 713 | 0,0237 | 0,558 | 41,2 | 99 | KF186567.1 |
| MERS hCoV 2c England-Qatar/2012 | 30112 | 714 | 0,0237 | 0,559 | 41,2 | 99 | KC667074.1 |
| MERS hCoV KNIH/002 05 2015 | 30108 | 711 | 0,0236 | 0,558 | 41,1 | 99 | MK796425.1 |
| camel MERS CoV Egypt/Camel/AHRI-FAO-1/2018 | 30106 | 722 | 0,0240 | 0,566 | 41,2 | 99 | MK967708.1 |
| camel MERS CoV UAE B73 2015 | 30123 | 716 | 0,0238 | 0,561 | 41,2 | 99 | MF598663.1 |
| camel MERS CoV Qatar 2 2014 | 30117 | 713 | 0,0237 | 0,559 | 41,2 | 99 | KJ650098.1 |
| camel MERS CoV UAE B39 2015 | 30123 | 711 | 0,0236 | 0,560 | 41,1 | 99 | MF598631.1 |
| bat CoV PREDICT/PDF-2180 | 29642 | 659 | 0,0222 | 0,525 | 41,2 | 82 | NC 034440.1 |
| bat CoV Neoromicia/PML-PHE1/RSA/2011 | 30111 | 626 | 0,0208 | 0,517 | 40,1 | 85 | KC869678.4 |
| **OC43 hCoV ATCC VR-759** | **30741** | **485** | **0,0158** | **0,481** | **36,8** |  | **NC 006213.1** |
| OC43 hCoV 1908A/2010 | 30719 | 464 | 0,0151 | 0,462 | 36,7 | 99 | KF923886.1 |
| OC43 hCoV HK04-01 | 30710 | 471 | 0,0153 | 0,469 | 36,7 | 99 | JN129834.1 |
| OC43 hCoV HK04-02 | 30722 | 469 | 0,0153 | 0,468 | 36,7 | 99 | JN129835.1 |
| OC43 hCoV MY-U1024/12 | 30716 | 463 | 0,0151 | 0,463 | 36,6 | 99 | KX538975 |
| OC43 hCoV GZYF-26 | 30530 | 457 | 0,0150 | 0,460 | 36,6 | 98 | MG197715 |
| bovine CoV Kakegawa | 31038 | 485 | 0,0156 | 0,471 | 37,0 | 96 | AB354579.1 |
| bovine CoV BCoV-ENT | 31028 | 498 | 0,0161 | 0,481 | 37,1 | 95 | NC 003045.1 |
| bovine CoV Mebus | 31032 | 484 | 0,0156 | 0,470 | 37,0 | 96 | U00735.2 |
| bovine CoV DB2 | 31007 | 486 | 0,0157 | 0,469 | 37,1 | 96 | DQ811784.2 |
| porcine CoV PHEV CC14 | 30682 | 504 | 0,0164 | 0,486 | 37,3 | 93 | MF083115.1 |
| porcine CoV PHEV USA-15TOSU1362 | 30515 | 506 | 0,0166 | 0,495 | 37,1 | 92 | KY419110.1 |
| porcine CoV PHEV VW572 | 30480 | 504 | 0,0165 | 0,490 | 37,2 | 93 | DQ011855.1 |
| **HKU1 hCoV** | **29926** | **340** | **0,0114** | **0,458** | **32,1** |  | **NC 006577.2** |
| HKU1 hCoV N25 genotype B | 29845 | 324 | 0,0109 | 0,440 | 32,0 | 95 | DQ415902.1 |
| HKU1 hCoV N22 genotype C | 29905 | 346 | 0,0116 | 0,468 | 32,0 | 96 | DQ415899.1 |
| HKU1 hCoV BJ01-p3 | 29887 | 338 | 0,0113 | 0,459 | 32,0 | 99 | KT779555.1 |
| rodent CoV RtNn-CoV/SAX2015 | 31172 | 585 | 0,0188 | 0,523 | 38,4 | 74 | KY370049.1 |
| rodent CoV<br>VZ BetaCoV 16715_52 | 31083 | 572 | 0,0184 | 0,530 | 37,8 | 74 | MH687968.1 |
| rodent CoV RtBi-CoV/FJ2015 | 31149 | 601 | 0,0193 | 0,533 | 38,6 | 74 | KY370051.1 |
| **229E hCoV** | **27317** | **488** | **0,0179** | **0,496** | **38,3** |  | **NC 002645.1** |
| 229E hCoV<br>229E/Seattle/USA/SC399/2016 | 27055 | 473 | 0,0175 | 0,492 | 38,0 | 97 | KY674914.1 |
| 229E hCoV 229E/Haiti-1/2016 | 27271 | 478 | 0,0175 | 0,494 | 38,0 | 98 | MF542265.1 |
| 229E hCoV 229E/BN1/GER/2015 | 27022 | 473 | 0,0175 | 0,490 | 38,1 | 97 | KU291448.1 |
| camel CoV<br>camel/Jeddah/N60/2014 | 27390 | 497 | 0,0181 | 0,499 | 38,4 | 92 | KT368899.1 |
| camel CoV camel/Taif/T96/2015 | 27392 | 498 | 0,0182 | 0,501 | 38,4 | 92 | KT368914.1 |
| camel CoV camel229E-CoV/JC49/KSA/2014 | 27193 | 490 | 0,0180 | 0,496 | 38,4 | 91 | KT253325.1 |
| bat CoV BtCoV/KW2E-F151/Hip cf. rub/GHA/2011 | 28026 | 519 | 0,0185 | 0,497 | 38,9 | 88 | KT253269.1 |
| bat CoV BtKY229E-8 | 27636 | 522 | 0,0189 | 0,502 | 39,1 | 85 | KY073748.1 |
| **NL63 hCoV Amsterdam I** | **27553** | **332** | **0,0120** | **0,417** | **34,5** |  | **NC 005831.2** |
| NL63 hCoV<br>NL63/Seattle/USA/SC0768/2019 | 27537 | 327 | 0,0119 | 0,419 | 34,1 | 98 | MN306040.1 |
| NL63 hCoV Amsterdam 057 | 27550 | 329 | 0,0119 | 0,412 | 34,5 | 99 | DQ445911.1 |
| NL63 hCoV Kilifi_HH_5402_20-May-2010 | 27832 | 327 | 0,0117 | 0,406 | 34,5 | 98 | MG428704.1 |
| bat CoV BtKYNL63-9a | 28363 | 609 | 0,0215 | 0,563 | 39,2 | 76 | KY073744.1 |

## Notes

Conflict of interest: The authors declare that no competing interests exist.

### Competing Interest Statement

The authors have declared no competing interest.

